# Zoledronic acid targets the mevalonate pathway causing reduced cell recruitment and attenuation of pulmonary fibrosis

**DOI:** 10.1101/2021.12.01.470755

**Authors:** Lloyd Tanner, Jesper Bergwik, Ravi KV Bhongir, Arne Egesten

**Author notes:** Corresponding author: Lloyd Tanner.

## Abstract

Idiopathic pulmonary fibrosis (IPF) is a progressive lung disease causing irreparable scarring of lung tissue, resulting in most patients succumbing rapidly after diagnosis. With limited treatment options available, repurposing of current pharmaceuticals offers an expeditious option to address this dire need. The mevalonate pathway, which is involved in the regulation of cell proliferation, survival and motility, is targeted by the bisphosphonate zoledronic acid (ZA). In this study, administration of ZA reduced myofibroblast transition and blocked NF-kB signaling in macrophages leading to impaired immune cell recruitment. ZA treatment of mice with bleomycin-induced lung damage displayed decreased levels of cytokines in the BALF, plasma, and lung tissue, resulting in less histologically visible fibrotic scarring. Additionally, bleomycin induced production of the ZA target, farnesyl diphosphate synthase (FDPS), was reduced in lung tissue and fibroblasts upon ZA treatment. Therefore, ZA administration offers an expedient and efficacious treatment option against IPF in a clinical setting.

**Teaser:** Repurposing of zoledronic acid potentially offers a clinically viable treatment for idiopathic pulmonary fibrosis

## Introduction

Idiopathic pulmonary fibrosis (IPF) is a disorder characterized by progressive lung scarring with a median survival time of 3 years postdiagnosis (*1*–*3*). The disease is associated with increasing cough and dyspnea, affecting approximately 3 million people worldwide, with incidence strongly correlated with increasing age (*4, 5*). Characteristic features of IPF include epithelial injury (*6, 7*), myofibroblast transition, migration, and proliferation (*8, 9*), inflammatory cell recruitment including macrophages and neutrophils (*10*), and finally extracellular matrix deposition and tissue distortion (*11*).

Current IPF therapies focus on the inhibition of collagen deposition by blocking myofibroblast activation, with limited success in achieving overall IPF resolution necessitating the need for novel therapies (*5*). Drug repurposing allows for shortened preclinical and clinical trial periods from an estimated 10-12 years to 3-4 years, allowing patients to benefit from cheaper medications (*12*–*14*). Drugs that exhibit multiple target sites are often viewed negatively given their potential for unwanted side effects. However, this polypharmacology presents opportunities to treat other disorders, particularly those exhibiting complex pathologies such as IPF (*15, 16*).

Bisphosphonates are a group of drugs used for osteoporosis treatment directed at osteoclasts to reduce bone turnover (*17*). Osteoclasts originate from myeloid progenitors but can also be derived from differentiating macrophages (*18*). The similarity between the osteoclasts and macrophages results in bisphosphonates targeting macrophages simultaneously, which has been applied in cancer treatment targeting tumor associated macrophages (TAMs) (*19*). Bisphosphonate treatment triggers repolarization of TAMs from M2 to M1, resulting in increased tumor killing (*20*). In an IPF context, an increased M2 macrophage population has been found in the blood of IPF patients as well as in the lungs of bleomycin-treated mice (*21*). The profibrotic role of M2 macrophages is achieved by an increased secretion of TGF-β1 (*22*), resulting in the promotion of fibroblast to myofibroblast transition and production of profibrotic proteins (*23*).

The primary target of bisphosphonates is farnesyl diphosphate synthase (FDPS), which is a key enzyme in the mevalonate pathway (*24*). The mevalonate pathway is a central metabolic pathway involved in steroid synthesis, N-glycosylation, protein geranylgeranylation and protein farnesylation (*25*). Furthermore, involvement of the mevalonate pathway has been shown in several human diseases, including cardiovascular disease, bone disorders, and cancer (*26*–*28*). This has resulted in the development of a wide range of pharmaceutical compounds, such as statins and bisphosphonates, that target enzymes in the pathway (*17, 29*). Interestingly, increased mevalonate pathway activity results in polarization of macrophages to an M2 state (*30*), suggesting that blocking the mevalonate pathway with bisphosphonates represents a viable drug target for IPF.

The third-generation bisphosphonate, zoledronic acid (ZA) is particularly interesting in this context due to use in the clinic and acceptable safety profile (*31*). The repurposing of bisphosphonates for use in pulmonary fibrosis-related disease is a unique concept that is not currently described in the literature. If successfully applied in humans, the novel treatment strategies described in this study would decrease the rate of disease progression, allow patients to mediate disease symptoms and increase their quality of life as well as improve the life expectancy associated with IPF.

## Results

### ZA reduces fibroblast to myofibroblast transition and production of fibrotic proteins

To determine zoledronic acid (ZA) involvement in fibroblast to myofibroblast transition (FMT), human lung fibroblasts (HFL-1) were utilized in a transwell assay (Fig.1A). Following stimulation with TGF-β1 for 48h, cell transition through the transwell membrane was measured. ZA treatment (10 μM) significantly reduced myofibroblast transition (*P* <0.001). Additionally, staining of mouse embryonic fibroblast cells (MEF) cells treated with ZA revealed the production of less collagen, fibronectin, and α-smooth muscle actin than in the TGF-β1 control, as measured by immunofluorescence (Fig. 1B).

**Figure 1:**
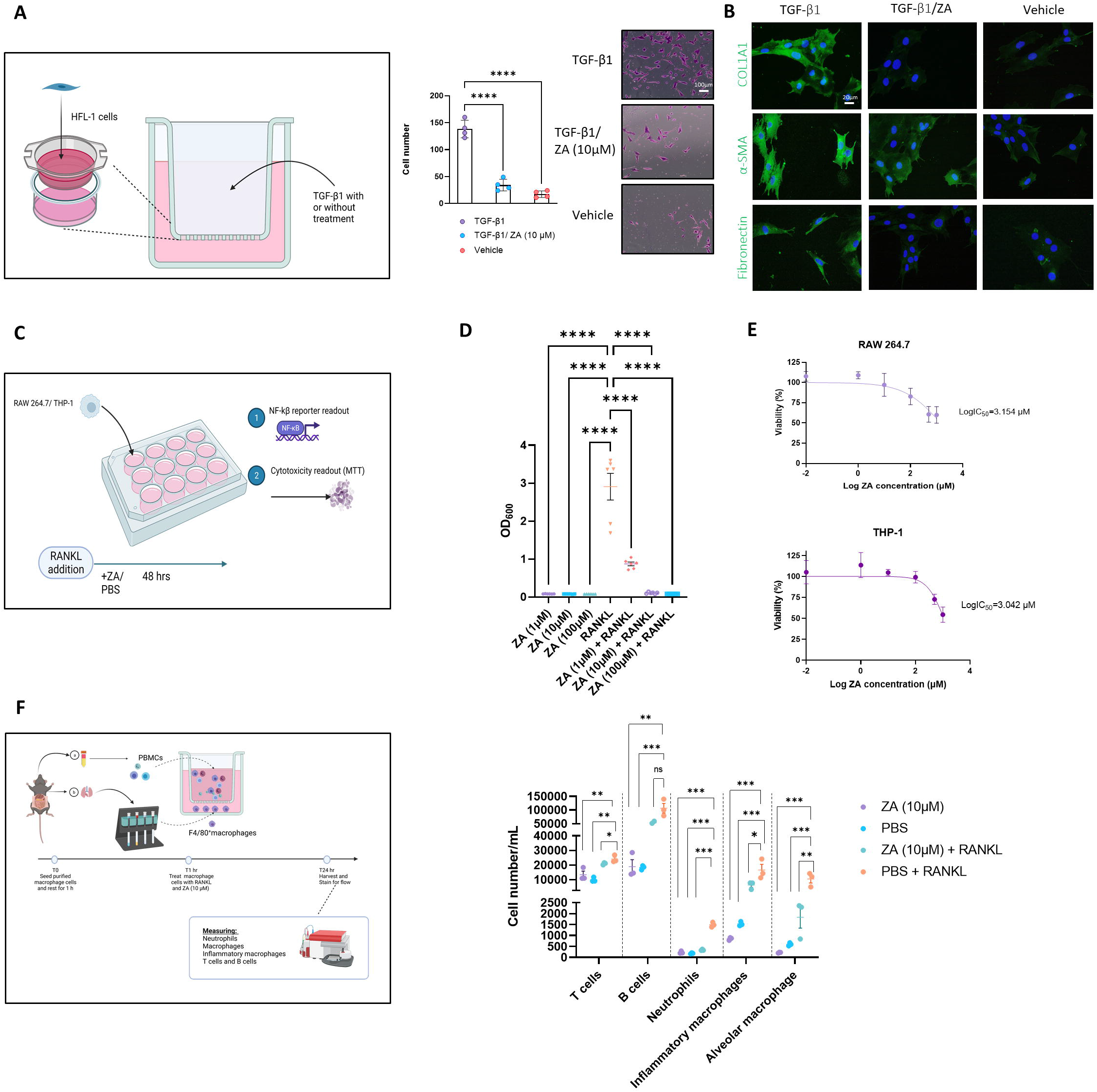
ZA inhibits *in vitro* transwell migration and fibrotic marker expression, whilst simultaneously reducing NF-κB signaling and immune cell recruitment. (A) ZA reduced myofibroblast migration and proliferation in a transwell assay compared to vehicle treatment and no treatment controls (Student’s *t*-test, *P<0*.*005* (***) and *P<0*.*0001* (****), with representative myofibroblast cells in lower chamber of transwell insert following 48h. (B) Immunostaining of fibroblast cells following 48h of TGF-β1 treatment, with ZA treatment displaying visually reduced levels of collagen, fibronectin and α-SMA compared to no treatment control. (C) RAW264.7 and THP1-Blue™ NF-κB cells were subjected to RANKL treatment for 48h, with (D) NF-κB signaling significantly reduced by ZA (*P*<0.0001) and (E) low levels of cytotoxicity reported for ZA against both THP-1 and RAW 264.7 cells. (F) Isolated murine macrophage cells (bottom well) and PBMCs (top well) were exposed to RANKL in a transwell system, with significant reductions in T cells, neutrophils, and macrophage cells following ZA treatment (Drug-treated samples were compared to untreated control samples using a one-way ANOVA followed by a Dunnett’s post-hoc test; *N*=3 technical replicates, from 3 biological replicates per measurement).

### Reduced NF-κB macrophage signaling moderates immune cell recruitment

NF-κB activation via canonical Wnt signaling in murine fibroblasts and myofibroblasts promotes the expression of tissue inhibitor of metalloproteinase 1 (TIMP1) and other pro-fibrotic factors in lung fibrosis (*32, 33*). To demonstrate the role of ZA in reducing NF-κB signaling, THP1-Blue™ NF-κB cells were stimulated with RANK ligand (RANKL) for 48h (Fig. 1C). ZA significantly reduced RANKL-induced NF-κB signaling at all tested concentrations (Fig. 1D) with ZA not displaying cytotoxicity in either RAW264.7 or THP1 cells at levels greater than Log 3 μM (Fig. 1E). Following reductions in NF-κB, we examined whether macrophage-mediated immune cell recruitment from peripheral blood was inhibited by ZA treatment. A wide variety of immune cells were measured using flow cytometry and revealed significant decreases between the RANKL and ZA/RANKL treated samples in recruitment of T cells, neutrophils, inflammatory macrophages, and alveolar macrophages (Fig. 1F).

### Zoledronic acid attenuates murine weight loss and fibrotic lung damage

Given the *in vitro* effect of ZA treatment in reducing fibrotic-related damage, a murine model of bleomycin-induced lung damage was employed, with ZA dosed either intratracheally (i.t.) or intraperitoneally (i.p.; 0.1 mg/kg) at day 15 (Fig. 2A). Murine weight loss was dramatically mitigated following i.p. ZA treatment, whilst i.t. ZA treatment offered less protection (Fig. 2B). The probability of survival was significantly increased following ZA i.p. administration (*P*<0.0329) whereas i.t. administration showed no statistically significant difference (Fig. 2C). Murine lung weight and hydroxyproline levels provide proxies for generalized lung damage and collagen content, with i.p. administration of ZA reducing both lung weight (*P*<0.0001) and hydroxyproline levels (*P*<0.0143) compared to vehicle/bleomycin mice (Fig. 2D and E). Images of murine lungs revealed decreased areas of fibrosis after i.p. treatment with ZA (Fig. 2F).

**Figure 2:**
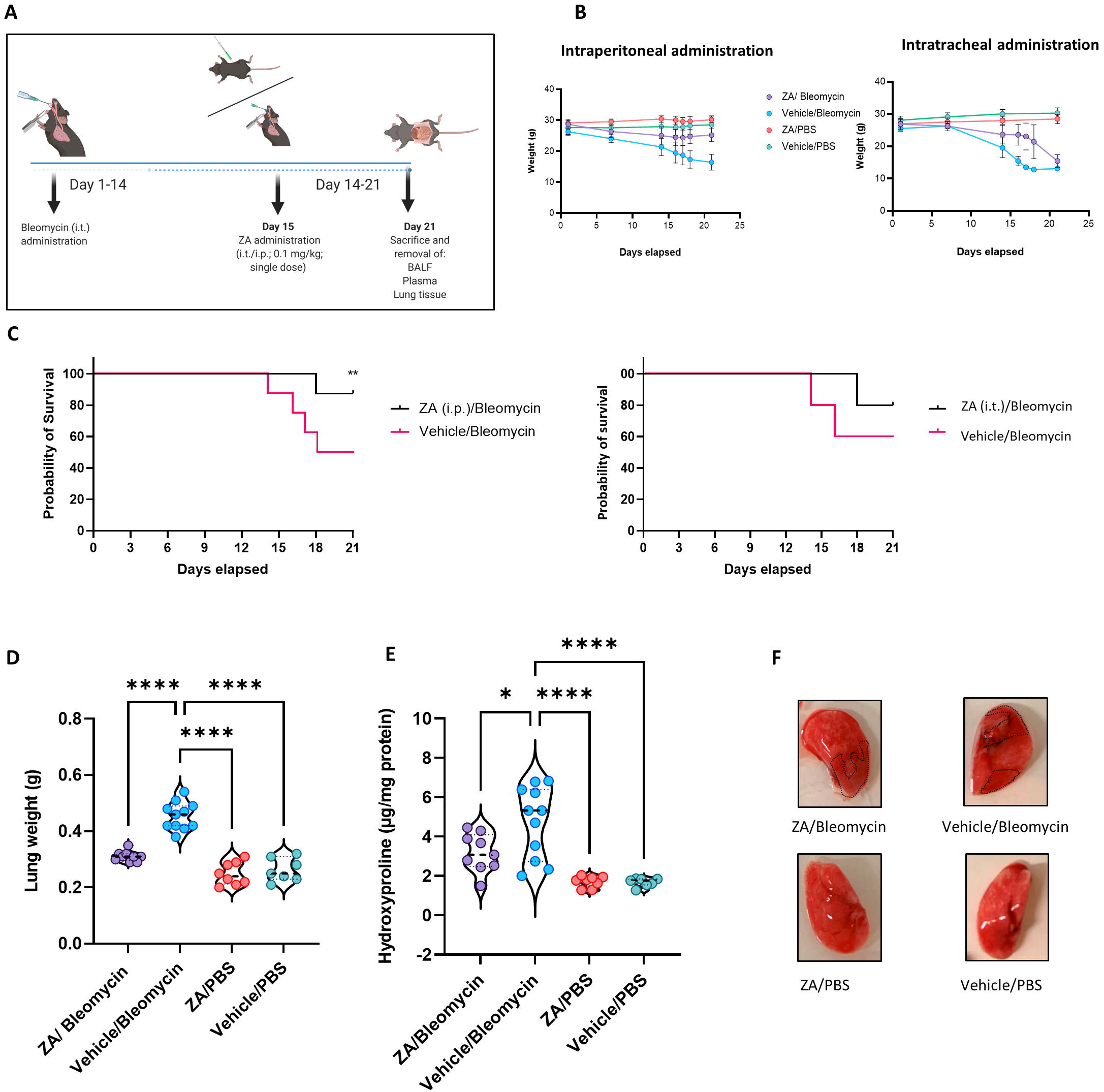
ZA administration, murine weights, survival curve, and lung readouts. (A) ZA was administered i.p. and i.t. on day 15 post-bleomycin introduction, with (B) murine total weight remaining stable in the ZA (i.p.) group for the duration of 21 days, with both bleomycin groups displaying significant weight loss after 14 days. (C) Resultant weight loss in the bleomycin/vehicle group caused significant mortality, with mice in the ZA/Bleo (i.p. and i.t.) groups seemingly protected. (D) Murine lung weight showed significant reductions in the ZA treated mice compared to the vehicle/bleomycin group (drug-treated samples were compared to untreated control samples using a one-way ANOVA followed by a Dunnett’s post-hoc test: \*\*\*\**P*<0.0001). Hydroxyproline collagen readout displayed significant reduction following ZA (i.p.) treatment (drug-treated samples were compared to untreated control samples using a one-way ANOVA followed by a Dunnett’s post-hoc test: \*\*\*\**P*<0.0001). (F) Representative murine images show bleomycin induced lung damage (highlighted in dotted areas).

### Murine immune cell recruitment to the lung is impaired by zoledronic acid

Administration of ZA effectively reduces regulatory T cell involvement in immune cell recruitment in cancer and fibrosis (*34, 35*), with a large proportion of these recruited cells being macrophages and neutrophils. Investigation of ZA treatment effects on immune cell infiltration into the airways was performed by flow cytometry on BALF obtained from *in vivo* bleomycin studies at Day 21. Decreases in neutrophil count and inflammatory macrophages were seen following both i.p. (*P*<0.0001) and i.t. (neutrophils: *P*=0.0417; inflammatory macrophages: *P*=0.0213) ZA administration compared to vehicle/bleomycin samples (Fig. 3A-C). Giemsa-Wright-stained BALF samples showed bleomycin-treated macrophages were enlarged and present in greater number, a phenomenon not seen in mice treated with ZA (i.p.; Fig. 3D).

**Figure 3:**
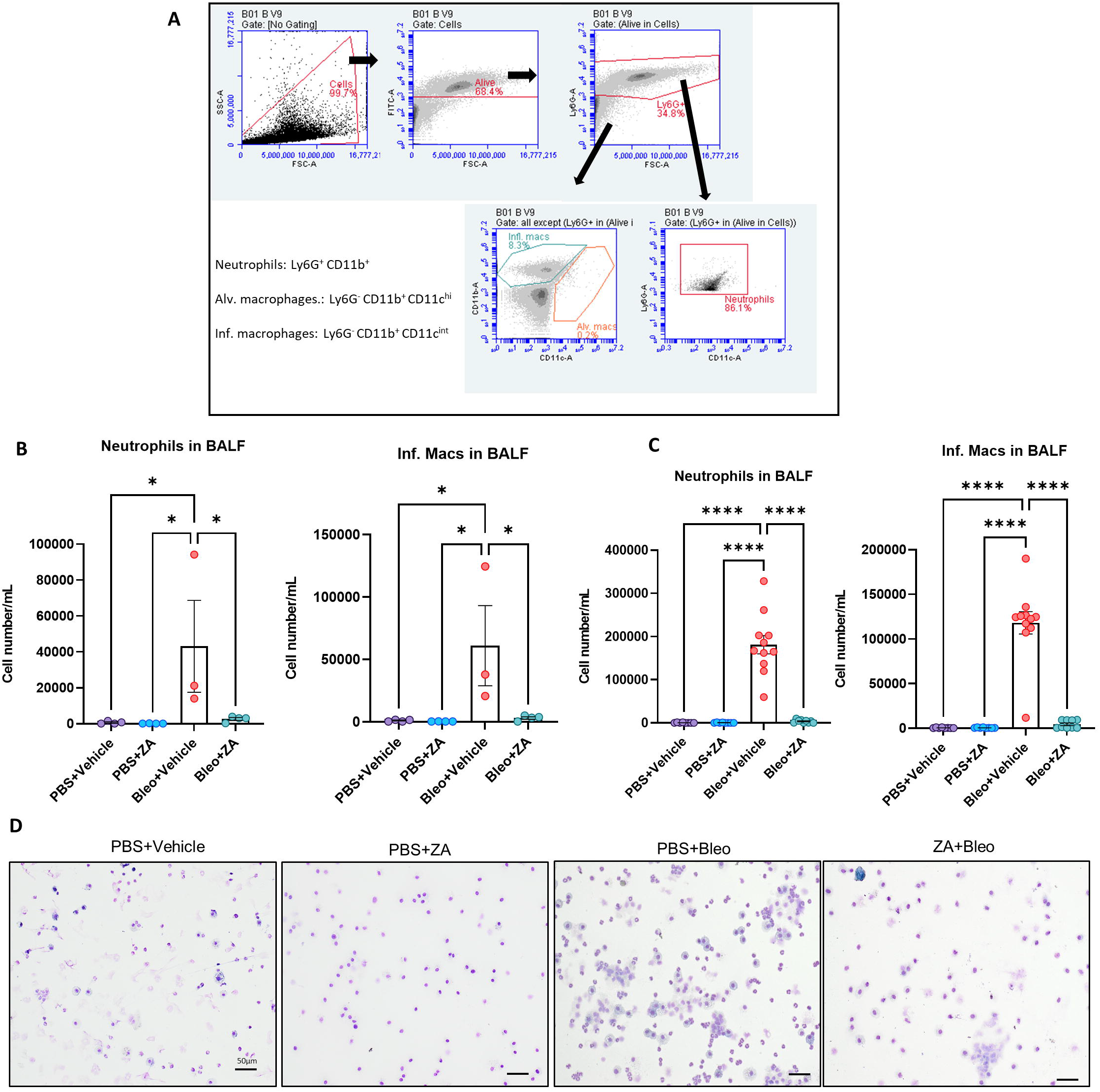
Inflammatory cell influx measured in murine BALF. Murine BALF was assessed for neutrophils and inflammatory macrophages (B and C) using flow cytometry, with the representative gating strategy depicted inset (A). Decreased numbers of neutrophils and inflammatory macrophages were detected in response to ZA (i.t.) treatment, with increased significant differences reported between the neutrophils and inflammatory macrophages of mice treated with ZA (i.p.). No significant differences were reported for alveolar macrophage numbers (not shown). Inflammatory cell numbers were compared to the vehicle/bleomycin group using a one-way ANOVA followed by a Dunnett’s post-hoc test (\**P*<0.05; *\*\*\*\*P*<0.0001). (D) Giemsa-Wright stained cytospin slides showing PBS/bleomycin BALF samples containing enlarged inflammatory macrophages, with ZA (i.p.) treatment reducing the presence of inflammatory macrophages. Scale bar=20 μm.

### Murine lung structure is maintained following intraperitoneal ZA administration

To determine whether murine histological damage was reduced more efficaciously by a single route of administration, both i.p. and i.t. lung samples from mice treated with ZA were assessed using H&E and picrosirius red staining (Fig. 4A and B; Supp. Fig. 1 and 2). ZA administered i.p. quantitatively reduced bleomycin-induced closure of the alveolar structures (*P*<0.0001), with significant decreases in collagen deposition (*P*<0.0001) seen in these samples compared to the vehicle/bleomycin samples. Conversely, intratracheally-administered ZA did not confer protection from lung damage as seen in both the H&E (*P*=0.7009) and picrosirius red staining (*P*=0.0839).

**Figure 4:**
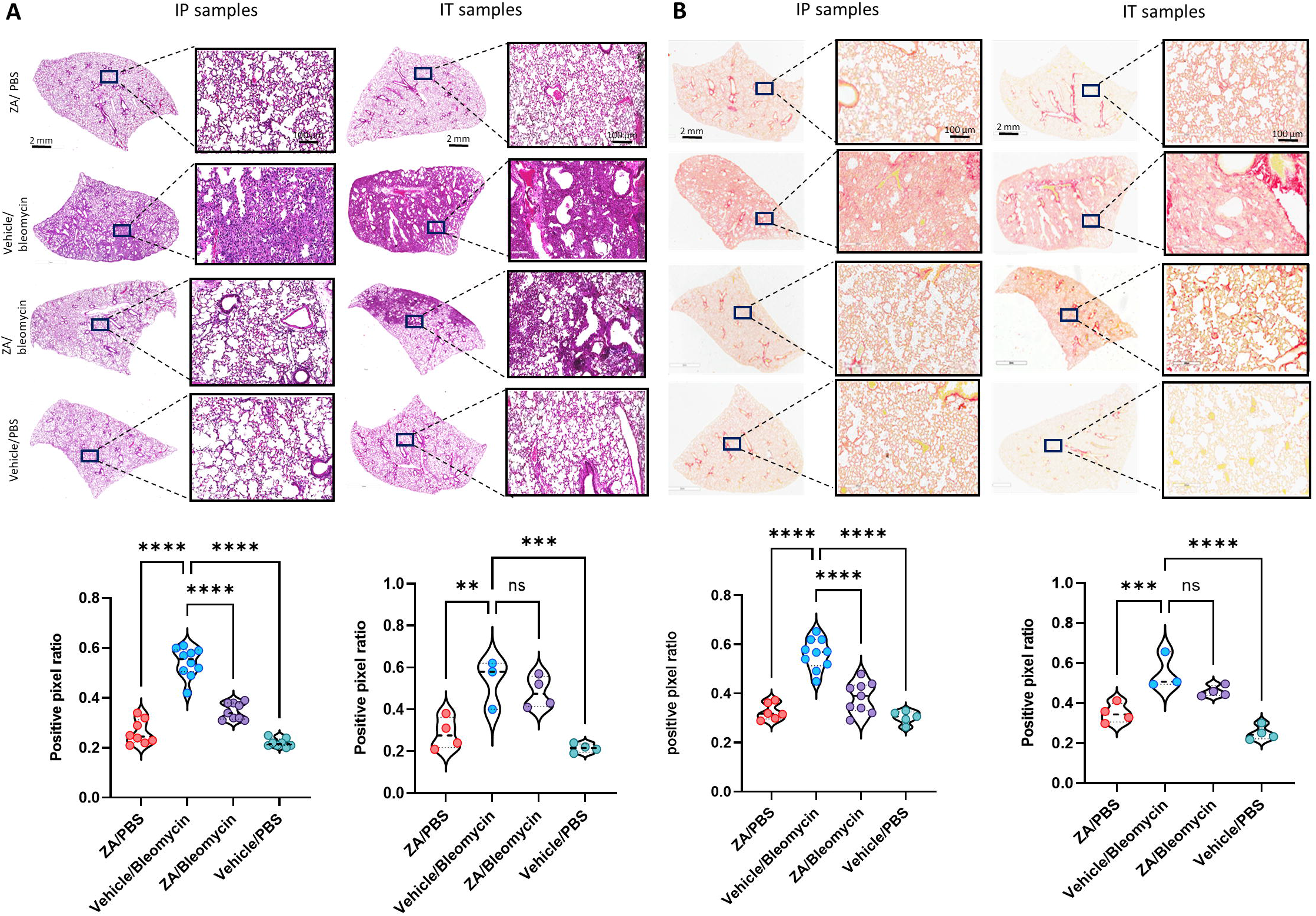
Murine lung staining displays reduced levels of fibrotic-related lung damage following ZA (i.p.) treatment compared to bleomycin/vehicle samples. (A) ZA (i.p.) significantly decreased lung damage in bleomycin-treated mice (H&E) and (B) collagen deposition (picrosirius red) in both macroscopic and microscopic structures compared to vehicle/bleomycin lungs and was confirmed by positive pixel analysis of whole-lung scanned images (scale bar of microscopic image=100 μm; scale bar of whole lung scan=2 mm). Statistical analyses were conducted using a one-way ANOVA followed by a Dunnett’s post-hoc test (*\*\*P<*0.01; *\*\*\*P*<0.005; \**P<*0.0001).

### Cytokine production elucidates decreased immune cell recruitment following ZA treatment

The effect of i.p. ZA treatment on inflammatory cytokine levels was investigated in murine samples from the *in* vivo study using a 23-cytokine inflammation multiplex assay (Supp. Fig. 3-5). The majority of cytokines were reduced in the BALF, plasma, and lung following ZA treatment. In particular, lung homogenate decreases were seen for IL-4 (*P*=0.0289), IL-6 (*P*=0.0007), GM-CSF (*P*<0.0001), KC (*P*<0.0001), and MCP-1 (*P*=0.0389) compared to the vehicle/bleomycin group (Fig. 5A-C). Decreased immune cell recruitment following ZA treatment in murine BALF is supported by reduced murine cytokine levels in murine BALF and plasma. Shared decreases were seen in both fluids for IL-6, GM-CSF, KC, MCP-1, and other cytokines indicative of reduced immune cell recruitment following ZA (i.p.) treatment (Fig. 5A and B; Supp. Fig. 3-5).

**Figure 5:**
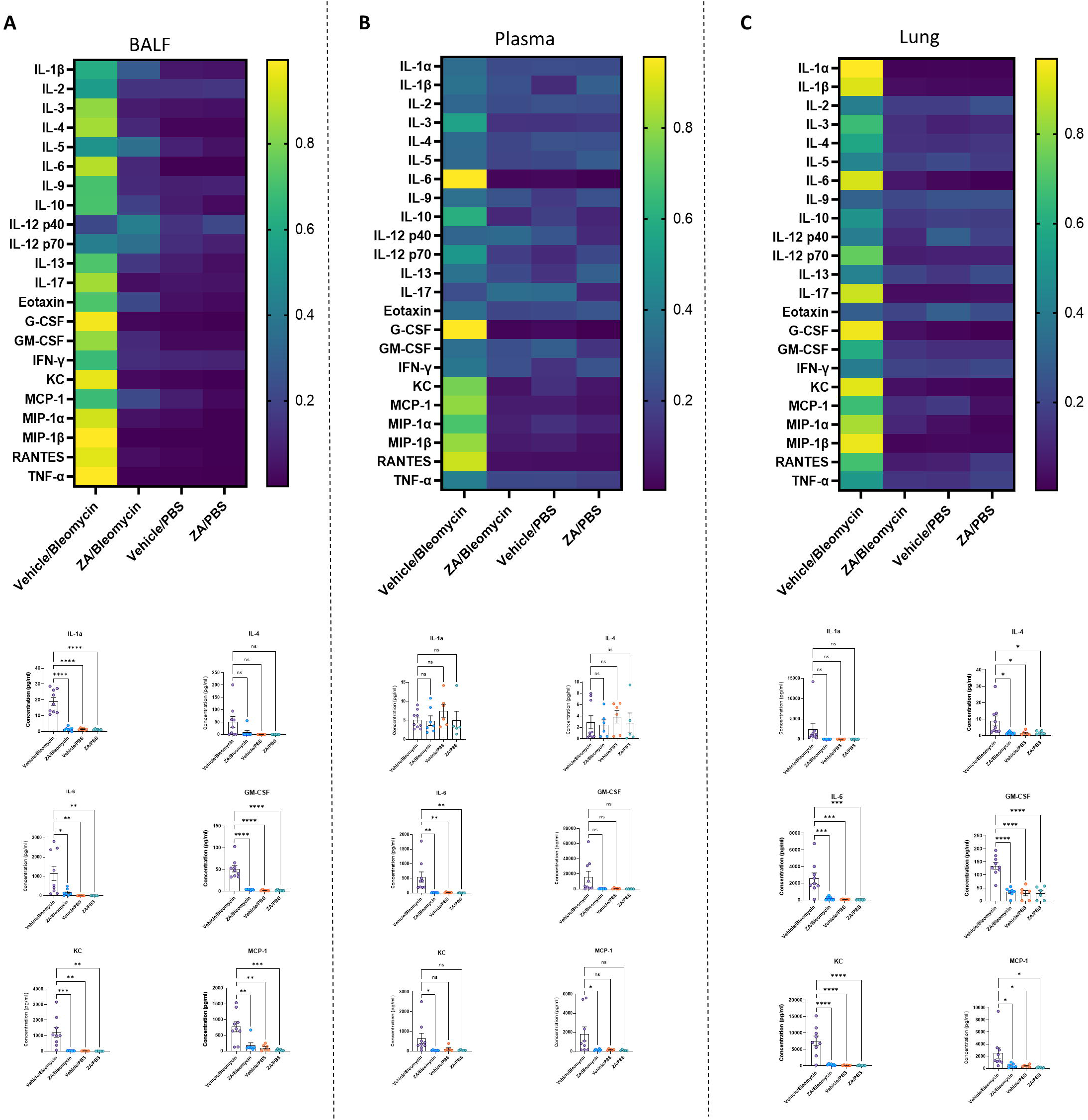
Significantly decreased murine cytokines following intraperitoneal ZA administration. Heatmaps (A-C) showing the differences in cytokine levels, as measured by multiplex assay, in murine BALF, plasma, and lung homogenate (yellow indicating high value; blue indicating low value) with selected corresponding significantly decreased or increased cytokine levels below. Cytokine values were compared to the vehicle/bleomycin group using a one-way ANOVA followed by a Dunnett’s post-hoc test (\**P*<0.05; *\*\*P*<0.01; *\*\*\*P*<0.005; *\*\*\*\*P*<0.001).

### Bleomycin-induced FDPS production in murine lung tissue is decreased by ZA treatment

Increased flux through the mevalonate pathway enhances posttranslational modification of Rac1, promoting macrophage/fibroblast signaling (*36*). Further elucidation of this mechanism was conducted by probing the downstream signaling pathway by assessing farnesyl diphosphate synthase (FDPS). Bleomycin-induced FDPS production was significantly decreased following ZA administration, with ZA/bleomycin samples displaying significantly reduced fluorescence (*P=*0.0077) and immunoreactivity compared to the vehicle/bleomycin group (Fig. 6A-C). Western blot analysis revealed reduced levels of FDPS in lung homogenates in the ZA/bleomycin group compared to the vehicle/bleomycin group (Fig. 6D). Furthermore, immunofluorescence staining of FDPS in mouse embryonic fibroblasts after TGF-β1 exposure showed an increase in FDPS, which was reduced upon ZA treatment (Fig. 6E).

**Figure 6:**
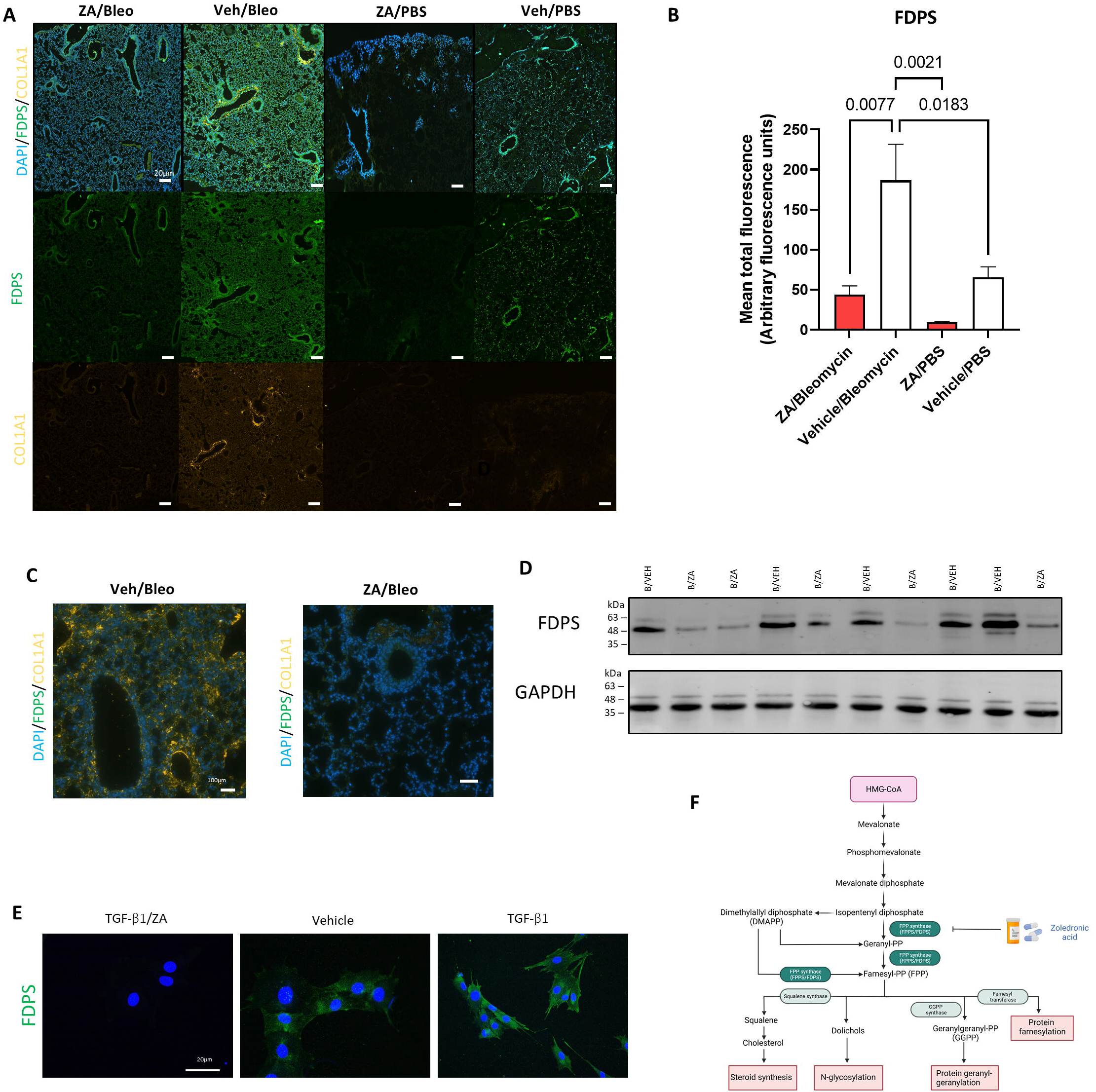
Lower production of FDPS and COL1A1 in murine lungs upon ZA treatment. (A) Murine lung immunofluorescence analysis showed decreased levels of immunoreactivity for FDPS and COL1A1 in ZA (i.p.)-treated mice compared to bleomycin (Bleo) controls (scale bar=20 μm), with (B) quantification of green fluorescence (ZA-treated and control samples were compared to bleomycin-treated samples using a one-way ANOVA followed by a Dunnett’s post-hoc test (\**P*<0.05; \*\**P*<0.01) and enlarged images of murine airways displaying co-staining of FDPS and COL1A1 (C). (D) Mouse lung homogenate samples analyzed by SDS-PAGE, followed by immunoblotting using rabbit antisera specific to FDPS revealed decreased protein levels within ZA-administered mice. (E) Murine embryonic fibroblast cells stimulated with TGF-β1 and treated with ZA/PBS for 48h displayed visually reduced levels of FDPS (scale bar=20 μm). (F) Mevalonate pathway depicting the main catalyzing enzymes and the target of ZA, FDPS.

## Discussion

IPF is characterized by progressive lung scarring, immune cell dysregulation, alveolar destruction, and eventual lung structure collapse (*5*). Theories surrounding IPF lung injury have focused on epithelial-derived causes of damage-associated molecular signaling without addressing critical myeloid-derived immune cell influence. With current drug targets and therapies predominantly directed toward fibroblast-mediated collagen deposition and fibrotic intercession, our focus turned to indirect intervention into immune cell-mediated inhibitory activities through mevalonate intermediate targeting. The mevalonate pathway has shown utility in multiple diseases, with evidence supporting mevalonate regulation of cell proliferation and energy homeostasis (*36*–*38*).

Statins have predominantly been used to probe the mevalonate pathway in interstitial lung diseases, offering mixed results. Whilst lung function decline and hospitalizations in IPF patients from the CAPACITY and ASCEND trials was diminished by statin use (*39, 40*), the post-hoc analysis of the INPULSIS trials did not establish beneficial effects of statin use in IPF patients (*41*). Furthermore, an FDA report associated statin use with onset of interstitial lung disease (*42*), with statins shown to increase lung fibrosis in a murine bleomycin model (*43*).

The approach reported in this study utilized a third-generation bisphosphonate, zoledronic acid (ZA), already employed for the treatment of osteoporosis, through potent anti-resorptive activity against osteoclasts (*44*). This activity is mediated by inhibiting farnesyl diphosphate synthase (FDPS), an enzyme responsible for catalysing the conversion of isopentenyl pyrophosphate and dimethylallyl pyrophosphate to their geranylated and farnesylated states, respectively (*45*). Herein, we report several arguments supporting FDPS targeting by ZA as an appealing and clinically relevant treatment for IPF.

Reported data indicate that increased geranylgeranylation substrates increase Rac1 activity in macrophages, with resultant polarization to a profibrotic phenotype (*36*). Rac1 also mediates ROS generation (*46*), with IPF-derived fibroblasts and macrophages generating higher levels of ROS (*47*). Furthermore, IPF-derived BAL cells displayed increased apoptotic resistance and increased TGF-β1 production (*48*). Our study found that TGF-β1-stimulated HFL-1 cells treated with ZA were less prone to myofibroblast transition and produced decreased levels of fibrotic proteins. RANKL induces phenotypic switching in macrophages from the initial state to M1-like cells, whereas IL-4, IL-10, IL-13, and other cytokines secreted by T helper cells are involved in M2 activation, skewing macrophages toward a profibrotic lineage (*49*). We showed that ZA-treated THP-1 cells stimulated with RANKL displayed reduced NF-κB signaling, a key regulator of proinflammatory responses in immune cells. In addition, *ex vivo* macrophage cells from murine BALF exposed to RANKL stimulation and ZA treatment displayed reduced cell recruitment efficiency.

Importantly, uncontrolled lung injury and immune cell recruitment is a hallmark of IPF initiation and progression, resulting in pro-inflammatory and pro-fibrotic cytokine release driving further fibrosis-related immune cell influx and ECM remodeling (*11, 50, 51*). Whilst recent efforts have focused on removal of senescent or damaged fibroblasts (*52, 53*), our approach advantageously suppresses pro-fibrotic mediators and immune cell recruitment with reduced cytotoxicity.

To elucidate the mechanism by which these effects were mediated, we examined murine lung samples. Whilst no direct targeting of FDPS has, to our knowledge, been reported, FDPS has shown involvement in the progression of cardiac remodelling, with FDPS-targeting siRNA’s attenuating murine cardiac fibrosis (*54*). This is supported in our study, with ZA’s ability to decrease FDPS production demonstrated in both murine lung tissue and directly in fibroblast cells, correlating with subsequent decreases in collagen deposition. In addition, our study examined the most appropriate route of ZA administration, a key factor in any drug discovery program. Surprisingly, intraperitoneal administration of ZA produced the most effective alleviation of lung fibrosis. We hypothesize that direct introduction of ZA to the murine lung may produce cytotoxic damage or, as seen in previous generations of bisphosphonate, result in elimination of the macrophage population entirely (*55*). Conversely, intraperitoneal introduction of ZA displayed the ability to repolarize murine macrophages to an anti-fibrotic phenotype (*56*).

Important limitations addressed in this study include, whether treatment using ZA can display utility in human IPF pathologies. Whilst the translational aspect of this study is suggested using murine lung sections, it is important to demonstrate that ZA treatment successfully reduces FDPS levels and subsequent IPF lung damage in human clinical trials. Transition to the clinical trials is eased by the pre-existing FDA approval of zoledronic acid (i.v.) for osteoporosis, multiple myeloma, and bone metastases in cancer.

Together, our findings demonstrate that zoledronic acid possesses a mechanism of action targeting FDPS to suppress pulmonary fibrosis, which is distinct from currently employed therapeutic interventions. This study highlights the downstream effects of FDPS inhibition, including decreasing myofibroblast transition, decreasing inflammatory cell recruitment, and eventual inhibition of fibrotic-related lung remodelling. These data support the use of ZA in clinical trials for the treatment of IPF.

## Materials and Methods

### Study design

The goal of this study was to test a zoledronic acid repurposing approach to inhibit FDPS, ultimately leading to the inhibition of fibrosis-related progression. Initial *in vitro* experiments in both murine macrophages and fibroblast cells displayed decreased fibrosis-related phenotypic features. Subsequent *in vivo* murine studies using intratracheally-administered bleomycin were chosen as well-established and relevant models of experimental lung fibrosis. Sample sizes were calculated by power analysis based on previous experience, feasibility, and to conform to the ARRIVE guidelines (arriveguidelines.org). For lung fibrosis experiments testing, *n* ≥ 4 to 11 mice per group were used to achieve statistical significance. Mice were randomly assigned to treatment groups, with sample sizes decided before commencing experiments. Downstream analyses were conducted with the investigator blinded to the treatment groups, and no animals were excluded as outliers from the reported dataset. All *in vitro* and *in vivo* experiments were performed in 2 to 4 technical replicates.

### Ethical approval

All animal experiments were approved by the Malmö□Lund Animal Care Ethics Committee (M17009-18).

### Fibroblast transwell experiment

Human lung fibroblasts (HFL-1; ATCC, Manassas, VA, USA) were cultured until confluent in complete growth medium supplemented with L-glutamine, 100U/mL penicillin, 100 µg/mL streptomycin, and 10% FBS. Fibroblast chemotaxis was measured using 24-well Nunc (8-µm pore size) Transwell inserts (Thermo Fisher, Waltham, MA, USA). Following confluence, cells were seeded (5 × 10^5^ cells/ml) into the upper chamber in FBS-free medium, whilst the lower chamber contained complete medium with additional 10 % FBS as a chemoattractant. Experiments were carried out in experimental triplicate, with each experiment containing 4 biological replicates. Medium containing TGF-β1 (10 mM) followed by the addition of medium containing ZA (10μM), medium only, or vehicle only (DMSO 2% and PBS pH 7.4). Following 48h, medium was removed and cells in the lower chamber were stained (crystal violet) and imaged using a Nikon Eclipse 80i Compound Fluorescent Microscope and analyzed using Nikon NIS Elements F4.60.00.

### Immunostaining of embryonic mouse fibroblasts

Murine C57BL/6 embryonic fibroblasts (MEFs) were seeded (1 × 10^4^ cells/mL) into 24-well plates containing rounded glass cover slips. TGF-β1 (10 µM) was added to each well, followed by the addition of medium containing ZA or vehicle only. Following 48h of treatment, cells were washed and fixed with ice cold methanol, containing 0.5% Triton-X100. Slides were blocked using Dako Protein Block (Agilent, Santa Clara, CA, USA) for 1h at room temperature and then stained with primary antibodies (overnight), rabbit anti-COL1AI, rabbit anti-αSMA, and rabbit anti-fibronectin (Abcam, Cambridge, UK). Alexa Fluor 488-conjugated goat anti-rabbit antibody (Invitrogen, Carlsbad, CA, USA) was used as secondary antibody (2h). Glass cover slips were removed and mounted onto glass slides, with nuclei counterstained using DAPI containing fluoroshield (Abcam). Images were visualized using a Nikon Eclipse 80i Compound Fluorescent Microscope and analyzed using Nikon NIS Elements F4.60.00.

### NF-κB reporter assay

THP1-Blue™ NF-κB cells (derived from the human THP-1 monocyte cell line by stable integration of an NF-kB-inducible secreted embryonic alkaline phosphatase (SEAP) reporter gene (InvivoGen, San Diego, CA, USA; 2 × 10^5^ per well) were cultured in 96 well plate using RPMI containing 10 % heat inactivated FBS, 100 IU/mL of penicillin, and 100 μg/mL streptomycin. Cells were stimulated with PMA (100ng/mL) for 24 h, followed by RANKL (200 ng/mL) and ZA (10μM) treatment for 48h. After incubation, cells were centrifuged at 300 x g for 5 min, with supernatants used for the NF-κB activation assay. The assay was completed by adding Quanti-Blue™ (InvivoGen) to the supernatants, followed by incubation at 37 °C for 30 min. The absorbance was measured at 600 nm using a VICTOR 1420 Multilabel plate reader (PerkinElmer, Waltham, MA, USA).

### MTT assay

Mosmann’s MTT [3-(4,5-dimethylthiazol-2-yl)-2,5-diphenyltetrazolium bromide] assay was used to determine cell viability in both RAW264.7 and THP-1 cells. After the addition of the compounds at a starting concentration of 1000□μM (serial diluted to 0.1□μM), and incubation for 48h, each well received MTT at a concentration of 5□mg/ml in phosphate-buffered saline (PBS), with blank samples receiving only medium and MTT. Each compound concentration tested in this study was completed in triplicate. DMSO was added to each well, followed by plate shaking for 5□min to dissolve the formazan crystals. The absorbance of the formazan salt was measured at 540□nm by a VICTOR 1420 Multilabel plate reader. The following formula (Equation 1) was used to calculate the cell viability:

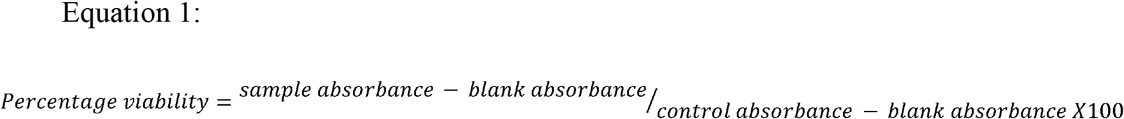

Nonlinear dose response curves were constructed using GraphPad Prism 4 software (GraphPad Software, San Diego, CA, USA) and Microsoft Excel.

### Ex vivo cell recruitment assay

BAL was performed on C57BL/6 mice (n=10) with a total volume of 1 ml PBS containing 100 μM EDTA. BALF was collected in Eppendorf tubes on ice and immediately separated using the Anti-F4/80 MicroBeads UltraPure (mouse; 130-110-443), dead cell removal kit (130-090-101), MS columns (130-042-201), and QuadroMACS™ Separator (130-090-976; Miltenyi Biotec, Bergisch Gladbach, Germany). Purified murine macrophages were counted and run on a BD Accuri C6 flow cytometer to determine cell purity. Concurrently, murine blood was isolated via heart puncture and collected in EDTA-containing tubes. Murine PBMCs were isolated using Ficoll-Paque PREMIUM 1.084 (Sigma-Aldrich, Saint Louis, MO, USA) density centrifugation (400 x *g* for 40 min) and subsequently counted. In the transwell plate, murine macrophages were added to the bottom well (5×10^5^ cells/well), with murine PBMCs (6×10^6^ cells/well) added to the top chamber. Following 24h of RANKL (200 ng/mL) macrophage stimulation, cells were harvested and stained for flow cytometry analysis carried out using a BD Accuri C6 (BD, Franklin Lakes, NJ, USA). Samples were separated into 3 aliquots, (i) alveolar macrophages as identified by F4/80 (BM8; 11-4321-42), CD11b (BD553312), and I-A/I-E (BD557000); (ii) B and T cells as identified by CD45 (BD553079), CD3 Molecular Complex (BD 555276), CD19 (BD561736); (iii) macrophage subtypes and neutrophils as identified by CD11b (BD553312), CD11c (BD558079), Ly-6G (BD551461).

### Animals

10-12-week-old male C57BL/6 mice (Janvier, Le Genest-Saint-Isle, France) were housed at least 2 weeks in the animal facility at the Biomedical Service Division at Lund University before initiating experiments and were provided with food and water *ad libitum* throughout the study. Mice were randomly allocated into five groups (n=4-11 animals per group): intratracheally (i.t.)-administered bleomycin (2.5 U/kg) + vehicle (i.p./i.t.), bleomycin (i.t.) +ZA (0.1 mg/kg; i.p./i.t.), saline (i.t.) + vehicle (i.p./i.t.), and saline (i.t.) + ZA (0.1 mg/kg; i.p./i.t.).

### Blood collection

Blood was collected in 0.5 M EDTA tubes by cardiac puncture and centrifuged at 1,000 x *g* for 10 min. Plasma supernatant was used for the analysis of inflammatory mediators using a multiplex assay (Bio-plex assay; Bio-Rad, Hercules, CA, USA).

### Collection of lung tissue

Right lungs were collected in Eppendorf tubes on dry ice and stored at -80 °C. The snap-frozen lungs were thawed and homogenized in tissue protein extraction reagent (T-PER) solution (ThermoFisher) containing protease inhibitor (Pefabloc SC; Sigma-Aldrich) at a final concentration of 1 mM. Lung homogenates were centrifuged at 9,000 x *g* for 10 min at 4°C, and the supernatants were collected for multiplex analysis. Left lungs were collected in Histofix (Histolab, Göteborg, Sweden) and submerged in 4 % buffered paraformaldehyde solution.

### Bronchoalveolar lavage fluid (BALF) collection

BAL was performed with a total volume of 1 ml PBS containing 100 μM EDTA. BALF was collected in Eppendorf tubes on ice, with aliquots made for flow cytometry, cytospin differential counts, and an aliquot transferred to -80°C for multiplex cytokine analysis. Cytospin preparations of cells were stained with modified Wright-Giemsa stain (Sigma-Aldrich).

### Flow cytometry

Flow cytometry was carried out using a BD Accuri C6 (BD). The washed cells were incubated with Fixable Viability Stain 510 (FVS510) (BD564406) to differentiate live and dead cells. Cells were washed with Stain buffer 1x (BD554656) and incubated with Lyse Fix 1x (BD558049 (5x)). Fixed cells were washed with stain buffer and aliquoted into two parts. One was incubated with CD11b (BD553312), CD11c (BD558079), Ly-6G (BD551461) antibodies and the other aliquot was incubated with CD11c, MHC (BD558593), SiglecF (BD562680) antibodies.

### Bioplex cytokine analysis

For the detection of multiple cytokines in BALF, plasma, and lung homogenate, the Bio-Plex Pro mouse cytokine assay (23-Plex Group I; Bio-Rad) was used on a Luminex-xMAP/Bio-Plex 200 System with Bio-Plex Manager 6.2 software (Bio-Rad). A cytometric magnetic bead-based assay was used to measure cytokine levels, according to the manufacturer’s instructions. The detection limits were as follows: Eotaxin (21372.02-1.15 pg/mL), GCSF (124018.4-6.97 pg/mL), GMCSF (1161.99-3.73), IFN-γ (14994.64-0.72 pg/mL), IL-1α (10337.5-0.63 pg/mL), IL-1β (28913.54-1.57 pg/mL), IL-2 (22304.34-1.21 pg/mL), IL-3 (7639.21-0.47 pg/mL), IL-4 (6334.86-0.36 pg/mL), IL-5 (12950.39-0.76 pg/mL), IL-6 (11370.16-0.66 pg/mL), IL-9 (2580.93-2.46 pg/mL), IL-10 (76949.87-4.09 pg/mL), IL-12p40 (323094.58-17.38 pg/mL), IL-12p70 (79308.46-19.51 pg/mL), IL-13 (257172.3-53.85 pg/mL), IL-17 (8355.61-0.5 pg/mL), KC (23377.88-1.3 pg/mL), MCP-1 (223776.6-45.04 pg/mL), MIP-1α (14038.07-0.58 pg/mL), MIP-1β (928.18-2.39 pg/mL), RANTES (4721.74-4.42 pg/mL), and TNF-α (73020.1-4.61 pg/mL). Cytokine measurements for samples were corrected for protein concentration, measured using a Pierce™ BCA Protein Assay Kit (Thermo Fischer Scientific).

### Hydroxyproline assay

Hydroxyproline levels in murine lung tissues were determined using the QuickZyme Hydroxyproline Assay kit (Quickzyme Biosciences, Leiden, The Netherlands). Lung tissues were homogenized as described above. Homogenates were diluted (1:1 vol:vol) with 12 N HCl and hydrolyzed at 95 °C for 20 h. After centrifugation at 13,000 × *g* for 10 min, 200 μl from the supernatant was taken and diluted 1:2 with 4 N HCl. Hydroxyproline standard (6.25–300 μM) was prepared in 4 N HCl and transferred to the microtiter plate. Following addition of a chloramine T-containing assay buffer, samples were oxidized for 20 min at RT. Detection reagent containing p-dimethylaminobenzaldehyde was added and after incubation at 60 °C for 1 h, absorbance was read at 570 nm with a VICTOR 1420 Multilabel plate reader (PerkinElmer, Waltham, MA, USA). The hydroxyproline content in lung tissue is given as hydroxyproline (μg) per mg lung tissue, corrected using a Pierce™ BCA Protein Assay Kit (Thermo Fisher Scientific).

### Immunostaining of murine lung sections

Lung tissue sections were fixed as reported above. Lung samples underwent antigen retrieval (pH 9 buffer) using a Dako PT Link pre-treatment module (Agilent). Samples were washed and blocked for 10 min (Dako protein block; Agilent, Santa Clara, CA) before being treated with primary antibodies overnight. Mouse anti-COL1A1 and rabbit anti-FDPS (Abcam, Cambridge, UK) antibodies were used. Alexa Fluor 488-conjugated goat/anti-mouse and Alexa Fluor 647 goat/anti-rabbit (Invitrogen, Waltham, CA, USA) were used as secondary antibodies. Glass cover slips were placed onto slides and mounted with DAPI-containing fluoroshield (Abcam). Microscopy was performed on a Widefield Epifluorescence Ti2 microscope equipped with a Nikon DS-Qi2 camera and fluorescence was quantified using ImageJ software.

### H&E and picrosirius red staining of lung tissue

A segment of the left lung was fixed in Histofix (Histolab) and embedded into paraffin and sectioned (3 µm) with a microtome. The tissue sections were placed on slides (Superfrost Plus; Fisher Scientific) and deparaffinized in serial baths of xylene and ethanol followed by staining using Mayer hematoxylin and 0.2% eosin (Histolab) or picrosirius red staining kit (Abcam, Cambridge, UK). The stained slides were imaged using an Olympus BX60F microscope with an SC50 camera.

### Statistical analysis

In this study, groups of three or more mice were compared using one-way analysis of variance (ANOVA) with Dunnett’s post hoc test. In experiments using two groups, results were compared using unpaired *t* test with Welch’s correction. Results in this study are displayed throughout as mean ± SEM. Statistical testing was carried out using GraphPad Prism 9.1.1 with statistical significance defined as *P* < 0.05.

## Acknowledgments

We would like to acknowledge the kind donation of MEF cells from Maria Allhorn (Lund University). We would also like to acknowledge the assistance received from the IQ Biotechnology Platform and the Lund University Bioimaging Centre (LBIC). In addition, we would like to acknowledge the flow cytometry support received from Assoc. Prof. Oonagh Shannon (Lund University). Figures feature elements created in BioRender.com.

## Funding

The work was supported by grants from:

Swedish Research Council 2020-011166 (AE)

The Swedish Heart and Lung Foundation 20190160 (AE)

The Swedish Government Funds for Clinical Research 46402 (ALF; AE)

The Alfred Österlund Foundation (AE)

## Data and materials availability

All data are available in the main text or the supplementary materials.

## Supplementary Materials

**Supplementary Figure 1: Whole lung scans of murine lungs from ZA experiments following H&E staining (scale bar=2 mm)**.

**Supplementary Figure 2: Whole lung scans of murine lungs from ZA experiments following picrosirius red staining (scale bar=2 mm)**.

**Supplementary Figure 3: Murine BALF cytokine levels from siRNA experiments**. Cytokine values were compared to the vehicle/bleomycin group using a one-way ANOVA (\**P<*0.05; *\*\*P<*0.01; *\*\*\*P<*0.005; *\*\*\*\*\*P<*0.001).

**Supplementary Figure 4: Murine plasma cytokine levels from siRNA experiments**. Cytokine values were compared to the vehicle/bleomycin group using a one-way ANOVA (\**P<*0.05; *\*\*P<*0.01; *\*\*\*P<*0.005; *\*\*\*\*\*P<*0.001).

**Supplementary Figure 5: Murine lung homogenate cytokine levels from siRNA experiments**. Cytokine values were compared to the vehicle/bleomycin group using a one-way ANOVA (\**P<*0.05; *\*\*P<*0.01; *\*\*\*P<*0.005; *\*\*\*\*\*P<*0.001).

