## Supplemental Figure 1-5 for "Zoledronic acid targets the mevalonate pathway causing reduced cell recruitment and attenuation of pulmonary fibrosis"

Short title: Repurposing zoledronic acid for pulmonary fibrosis

**Authors**

[Lloyd Tanner](https://www.sciencedirect.com/science/article/pii/S156919932030802X" \l "!)^1*^, Jesper Bergwik^1^, Ravi KV Bhongir^1^, Arne Egesten^1^

**Affiliations**

^1^Respiratory Medicine & Allergology, Department of Clinical Sciences Lund, Lund University and Skåne University Hospital, Lund, Sweden

**Supplemental Figures**

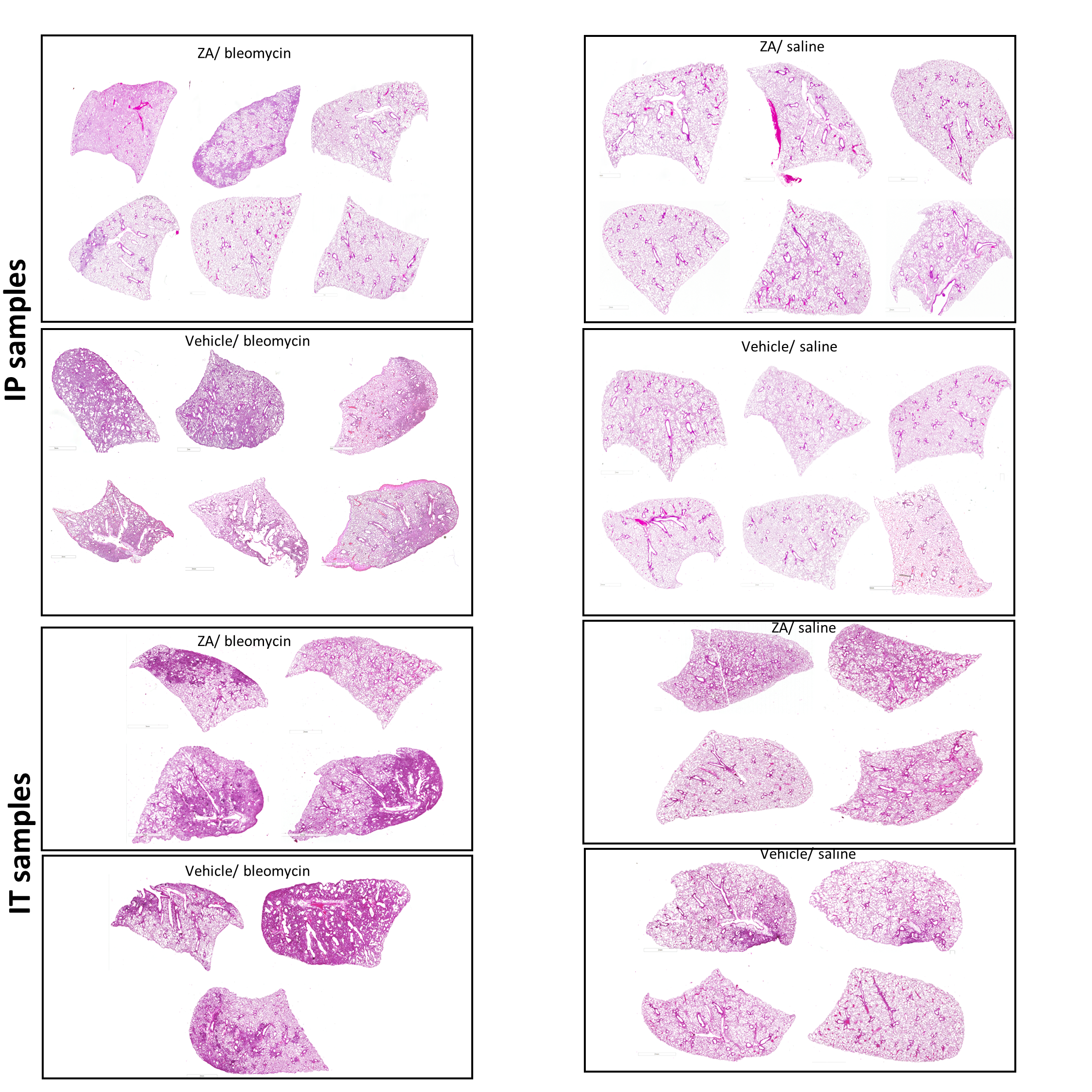

**Supplementary Figure 1: Whole lung scans of murine lungs from ZA experiments following H&E staining (scale bar=2 mm).**

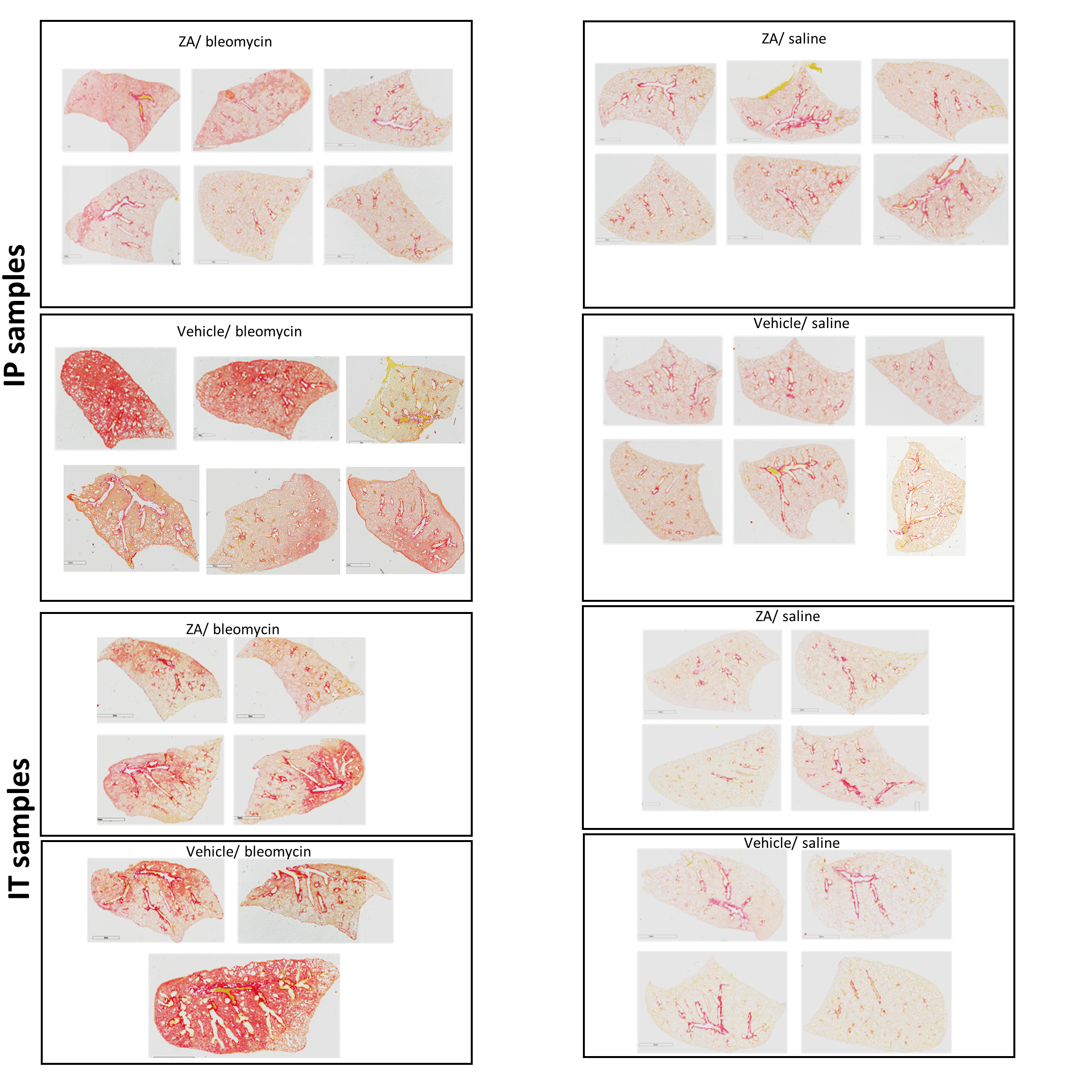

**Supplementary Figure 2: Whole lung scans of murine lungs from ZA experiments following picrosirius red staining (scale bar=2 mm).**

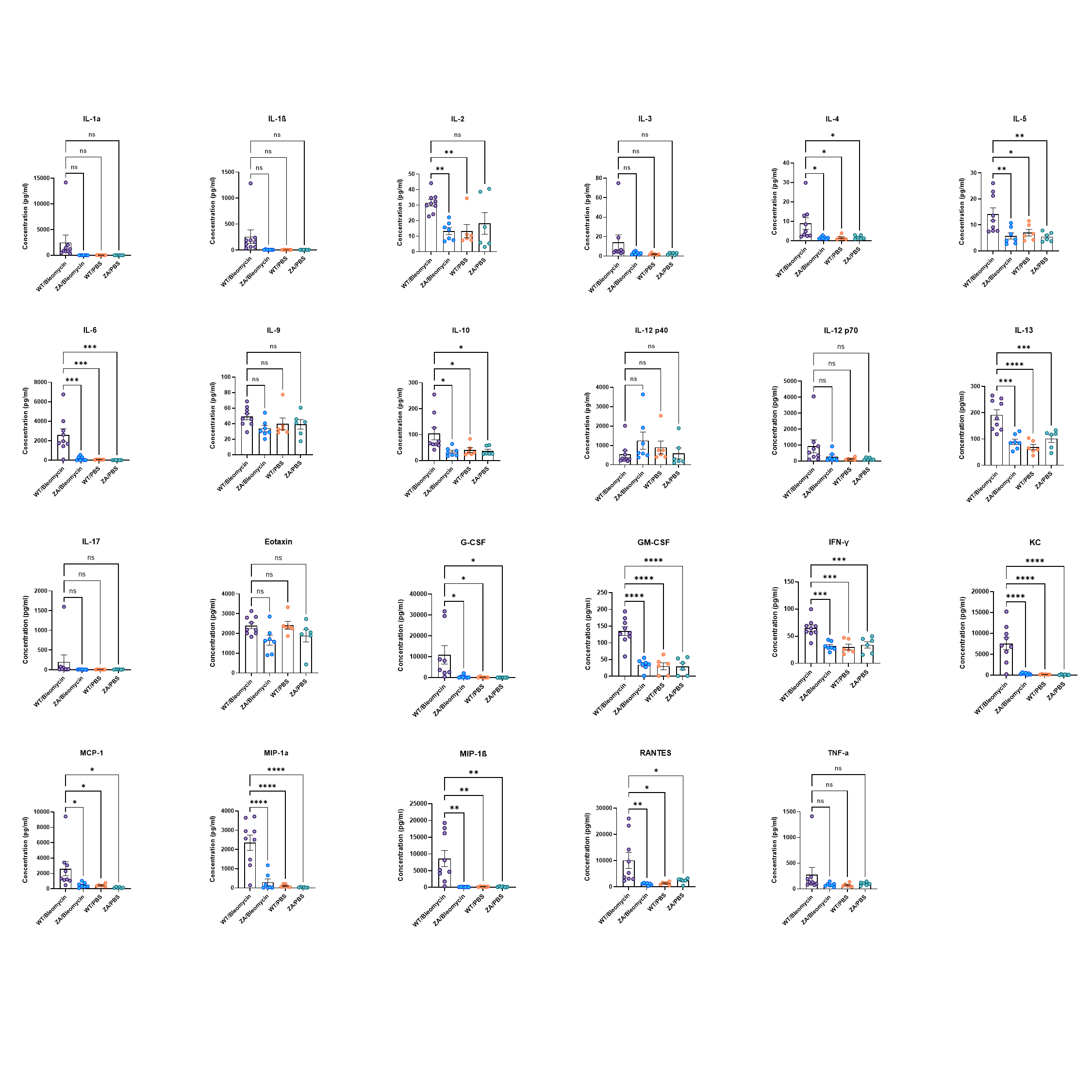

**Supplementary Figure 3: Murine BALF cytokine levels from ZA (i.p.) experiments.** Cytokine values were compared to the vehicle/bleomycin group using a one-way ANOVA (**P<*0.05*; **P<*0.01*; ***P<*0.005*; *****P<*0.001).

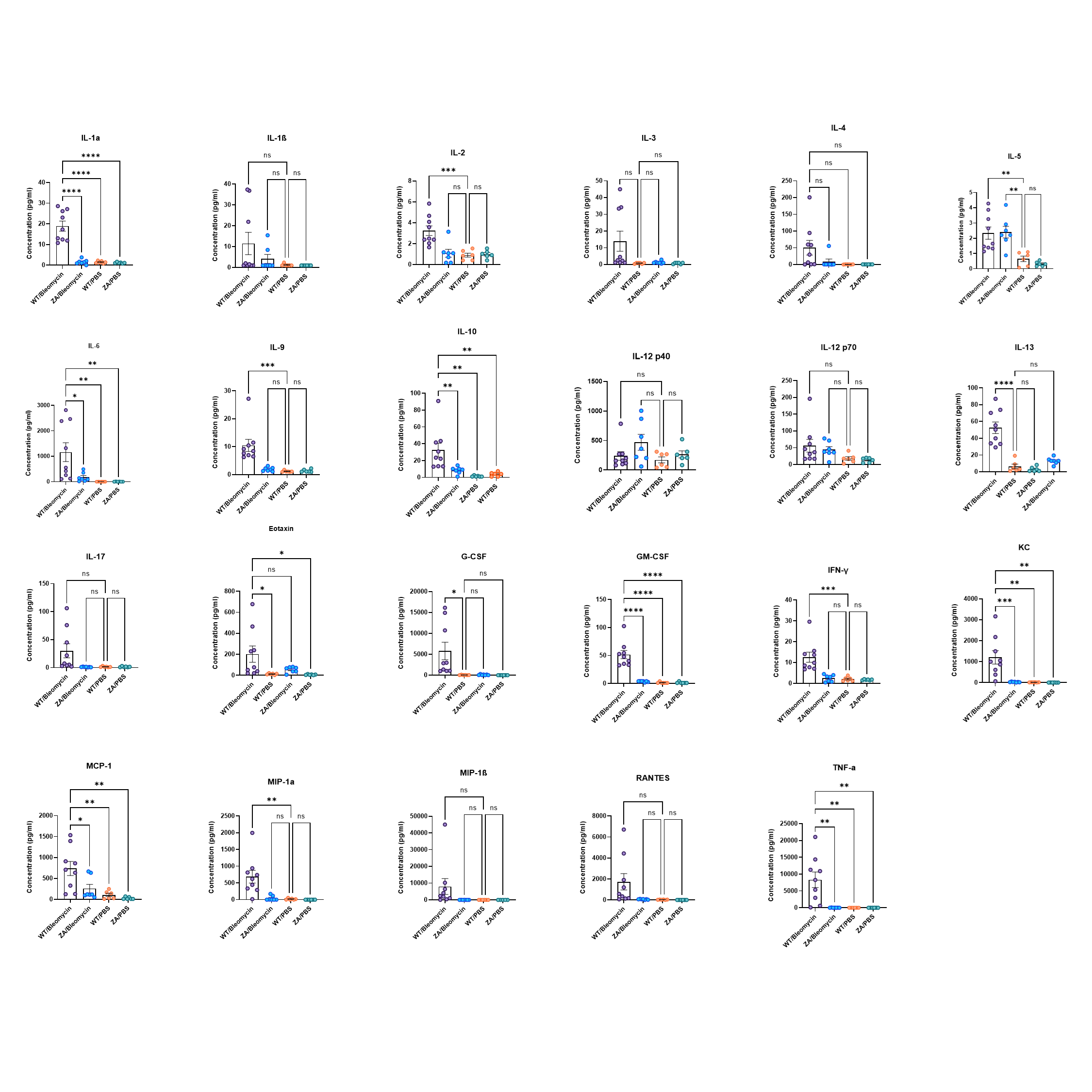

**Supplementary Figure 4: Murine plasma cytokine levels from ZA (i.p.) experiments.** Cytokine values were compared to the vehicle/bleomycin group using a one-way ANOVA (**P<*0.05*; **P<*0.01*; ***P<*0.005*; *****P<*0.001).

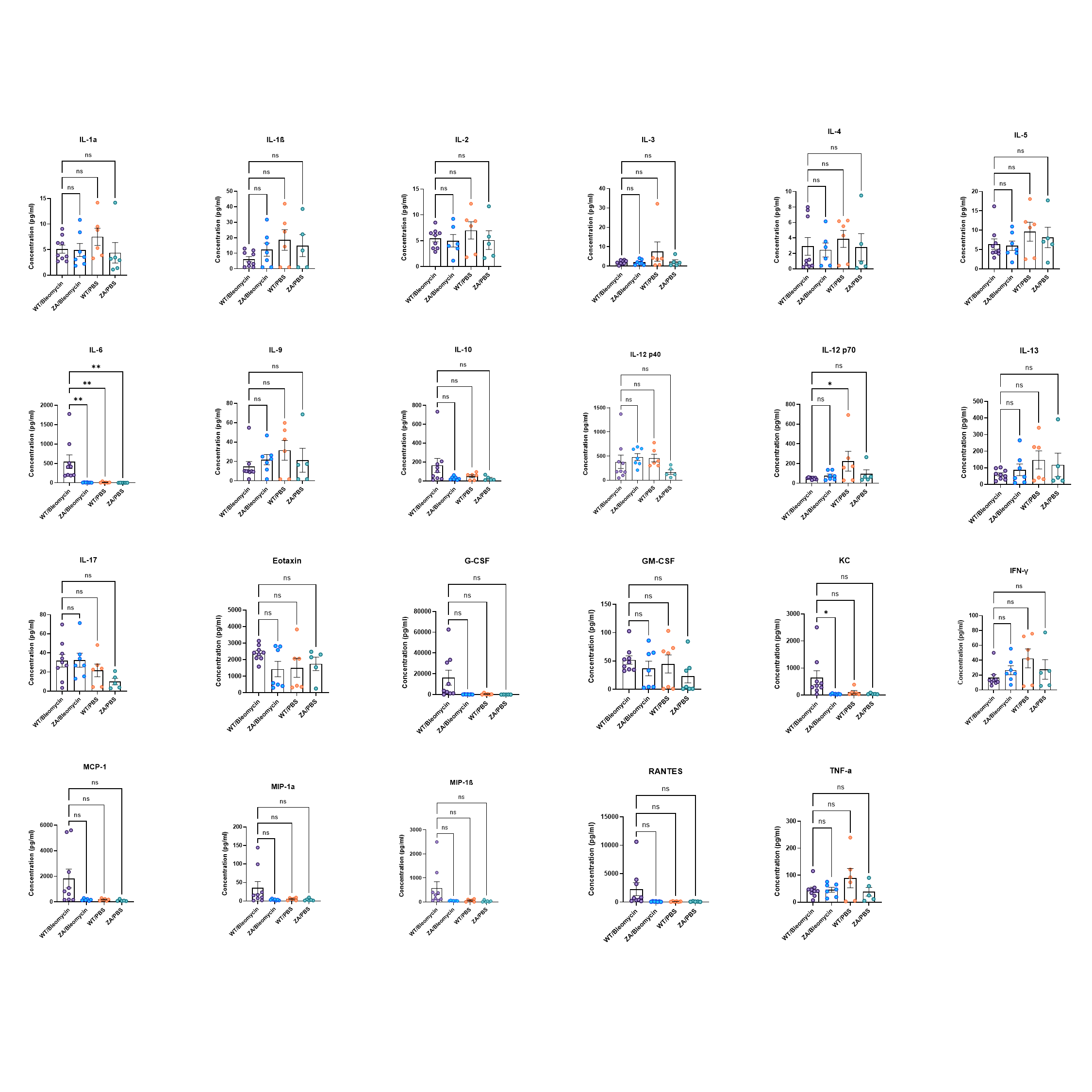

**Supplementary Figure 5: Murine lung homogenate cytokine levels from ZA (i.p.) experiments.** Cytokine values were compared to the vehicle/bleomycin group using a one-way ANOVA (**P<*0.05*; **P<*0.01*; ***P<*0.005*; *****P<*0.001).
